# Enhancement of brain atlases with laminar coordinate systems: Flatmaps and barrel column annotations

**DOI:** 10.1101/2023.08.24.554204

**Authors:** Sirio Bolaños-Puchet, Aleksandra Teska, Juan B. Hernando, Huanxiang Lu, Armando Romani, Felix Schürmann, Michael W. Reimann

**Author notes:** Co-lead authors (Sirio Bolaños-Puchet, Aleksandra Teska).

## Abstract

Digital brain atlases define a hierarchy of brain regions and their locations in three-dimensional Cartesian space. They provide a standard coordinate system in which diverse datasets can be integrated for visualization and analysis. Although this coordinate system has well-defined anatomical axes, it does not provide the best context to work with the complex geometries of layered brain regions such as the neocortex. To address that, we introduce *laminar coordinate systems* that consider the curvature and the laminar structure of the region of interest. These new coordinate systems consist of a principal axis, locally aligned to the vertical direction and measuring depth, and two other axes that describe a flatmap, a two-dimensional representation of the horizontal extents of layers. The main property of the flatmap is that it allows seamless mapping of information back and forth between 2D and 3D spaces, in a way consistent with the principal axis. It involves a structured dimensionality reduction where information is aggregated along depth. We propose a method to enhance brain atlases with laminar coordinate systems and flatmaps based on user specifications and define a set of metrics to characterize the quality of flatmaps. We applied our method to an atlas of rat somatosensory cortex based on Paxinos and Watson’s rat brain atlas, enhancing it with a laminar coordinate system adapted to the geometry of this region. Further, we applied our method to enhance the Allen Mouse Brain Atlas Common Coordinate Framework version 3 with a flatmap of the whole isocortex. We used this flatmap to produce new annotations of 33 individual barrels and barrel columns in the barrel cortex. Thanks to the properties of the flatmap, the resulting annotations are non-overlapping and follow the curvature of the cortex. Additionally, we introduced several applications highlighting the utility of laminar coordinate systems for data visualization and data-driven modeling. We provide a free software implementation of our methods for the benefit of the community.

## 1 Introduction

Experimental and computational studies in neuroscience make increasing use of standardized atlases describing the location and extents of brain structures in reference geometric spaces. These atlases are available for a variety of species, including rats (Paxinos and Watson, 2007; Papp et al., 2014), mice (Paxinos and Franklin, 2019; Wang et al., 2020) and humans (Amunts et al., 2020). By using registration methods, it is possible to place data from different origins and across different modalities into the common coordinate space of such an atlas. Data integrated in this way can be represented and analyzed in novel ways (Leergaard and Bjaalie, 2022), and provides the building blocks for data-driven modeling (Reimann, Bolaños-Puchet, et al., 2022).

While traditional printed atlases consist of a series of labeled outlines of brain slices along one or more cut planes, digital brain atlases represent information as a three-dimensional matrix where each voxel is labeled with the region to which it belongs. Typically, the Cartesian coordinate system defined by the voxel indices is aligned to match the anatomical axes of the brain (transverse, coronal, sagittal). As such, digital brain atlases provide a lookup for the locations of brain regions in global anatomical coordinates.

Although the Cartesian coordinate system is anatomically well-defined, it does not necessarily provide the best context in which to describe the shape and structure of particular brain regions. Thus, many regions have been traditionally analyzed and modeled with respect to region-specific coordinate systems that better capture their functional and developmental organization. For example, hippocampus has a complex geometry where a path following its local depth axis tends to be curved, and different paths can have entirely opposite orientations in the different hippocampal subfields (Strange et al., 2014). Since these properties cannot be accounted for by simple linear or affine transforms of the Cartesian coordinate system, hippocampal geometry has been described in terms of long-axis, proximal-distal and laminar coordinates in human (DeKraker et al., 2018) or longitudinal, transverse and radial coordinates in rat (Romani et al., 2023). Even thalamic nuclei contain functional gradients that can be expressed as axes of a region-specific coordinate system (Sumser et al., 2017).

Furthermore, as data is recorded over increasingly larger portions of the brain, the existence of continuous region-wide coordinate systems begins to constitute a requirement for data integration and analysis (L. Gao et al., 2022). Also, the construction of large-scale models of entire brain regions is likely to rely on algorithms that are defined in such coordinate systems adapted to the particular geometry of each region. Therefore, it becomes imperative to provide a reproducible and objective way to define these region-specific coordinate systems.

In the case of layered regions such as the neocortex, it is possible to define a particular kind of coordinate system that considers the curvature of the region as well as the laminar structure, providing a powerful tool to work with these complex geometries. We name this a *laminar coordinate system* and formalize its properties. First, the coordinate system is primarily defined by one *principal axis* of organization whose orientation varies smoothly in space. Second, the local orientation of this principal axis can be determined by the direction along which the layers change at each point in space. Third, the other two axes are locally orthogonal to the principal axis, but otherwise their orientation is of secondary importance.

The two axes orthogonal to the principal axis describe a *flatmap* of the region. The concept of a flatmap bears a resemblance to cartographic projections of the globe (Hahn and Duckworth, 2023), but is more complex, as it projects a three-dimensional structure and not just its surface. The main property of a flatmap is that it allows the seamless mapping of information back and forth between 2D and 3D spaces in a way consistent with the principal axis. It involves a kind of structured dimensionality reduction where information is aggregated in a direction orthogonal to the layers. For example, a circle drawn on the flatmap corresponds to a 3D subvolume similar to a cylinder, whose axis follows the local direction of the principal axis, and where the layer structure is preserved; in the context of the neocortex, such a subvolume would represent a cortical column, if appropriately sized. Conversely, any point inside this column would map back to the circle drawn on the flatmap.

In this manuscript, we introduce methods to define laminar coordinate systems and flatmaps based on user specifications, and provide a free software implementation for the benefit of the community. In order to ensure flexibility, and given the spatial discretization of the voxelized inputs, we do not attempt to derive a mathematical description of the laminar coordinate system. Instead, we represent it in the form of an *auxiliary atlas*, a set of three-dimensional matrices in the same Cartesian space as the base atlas, containing values for all coordinates at the center of each voxel. Additionally, we also provide the principal axis vector at each voxel. We call this process of generating auxiliary atlas datasets *atlas enhancement*. Our approach to provide not only a static flatmap, but tools to create and optimize them based on use case, sets us apart from previous work such as the flatmap of mouse isocortex in Wang et al., 2020.

As a result of the application of our methods, we enhance an atlas of rat somatosensory cortex (based on Paxinos and Watson, 2007) with a laminar coordinate system adapted to the shape of this region. We also enhance the Allen Mouse Common Coordinate Framework version 3 (CCFv3, Wang et al., 2020) by generating a flatmap of the whole mouse isocortex and using it to refine the annotations in the barrel cortex area (SSp-bfd) with individual barrels and associated barrel columns. Similar to a cartographic projection, a flatmap inevitably introduces some distortions. We therefore also define several metrics that can be used to quantify local and global distortions and assess the quality of the results. Finally, we present some applications of flatmaps for data-driven modeling and data analysis and visualization.

## 2 Methods

We introduce methods to generate the three axes of a laminar coordinate system in the form of auxiliary atlases. We describe the algorithms in detail but leave out the particularities of our implementation; these can be found in Supplementary material B. Specific values for the various parameters of the algorithms are provided in the Results section for the use cases presented there.

### 2.1 Generating depth and orientation fields

In order to define the principal axis, we propose an algorithm that generates auxiliary atlases of depth and local orientation. Specifically, depth information at each voxel is provided by a relative depth value (between 0 and 1) and a local thickness (in *µ*m); absolute depth (in *µ*m) can be easily computed as the product of these two. Local orientation is provided by a vector defined at each voxel; this vector is represented by a rotation of the Cartesian Y axis and stored as a rotation matrix.

The input to the algorithm is a voxelized brain atlas with region annotations, i.e., each voxel has a label of the brain region to which it belongs. We select a subset of regions of interest, e.g. all isocortex regions or somatosensory regions, and use them as the *source volume* for the following steps.

We begin by selecting top and bottom voxel shells in the source volume. These shells are located at the boundary of the source volume and have a thickness of one voxel; they represent zero depth and maximum depth, and would correspond to the cortical surface and the white matter boundary of the neocortex, respectively (Fig. 1A). Given the complexity of this task in actual brain region geometries (see Fig. 1D for an illustration in a simple geometry), it is performed by manual expert annotation. In short, we select all voxels that are in contact with the outside of the source volume, and then we remove all voxels on the sides (not at the top or bottom). The result of this process is an auxiliary atlas where each voxel is labeled as exterior, boundary (top, bottom or sides) or interior.

**Figure 1:**
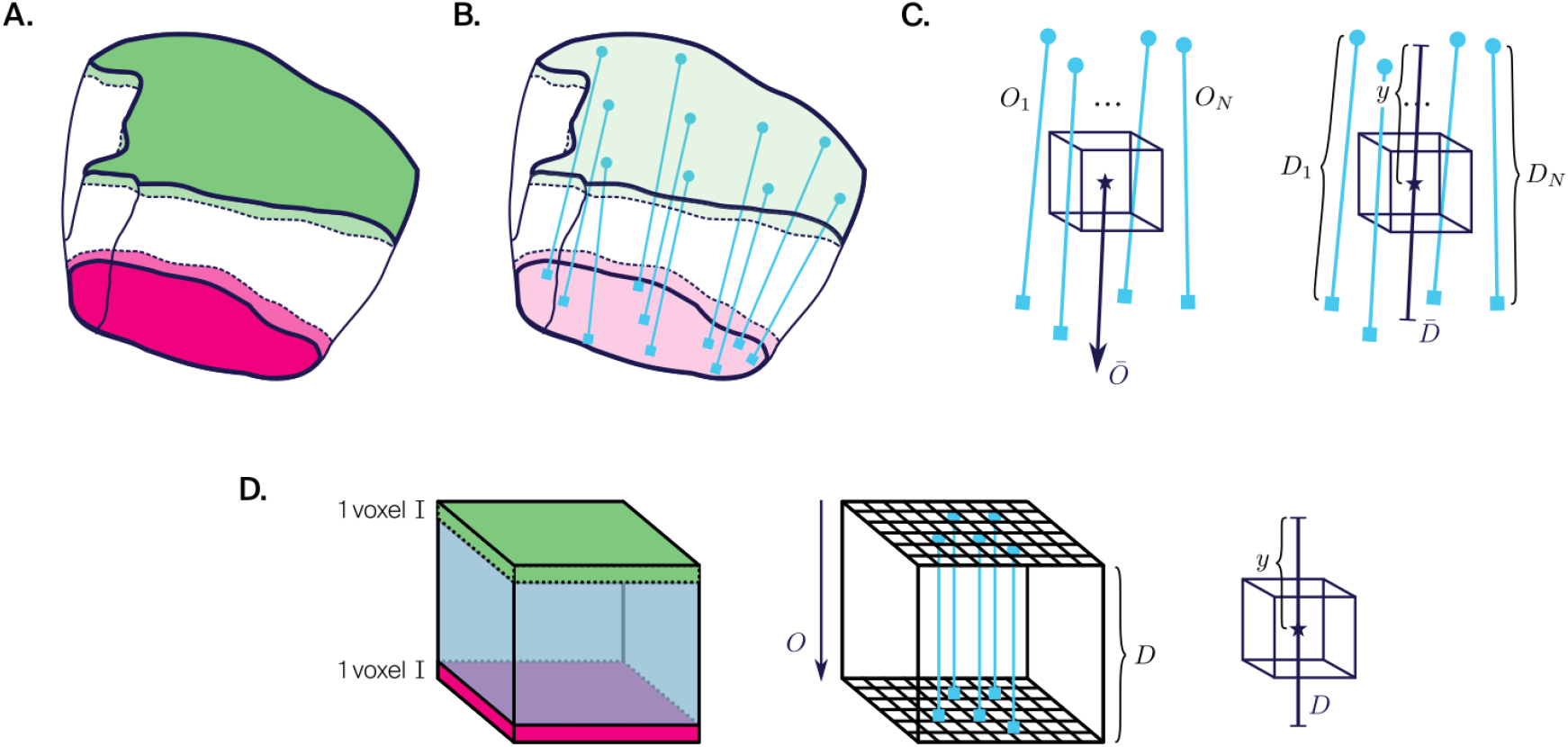
Generating depth and orientation fields. **A**. Top (green) and bottom (pink) shells. **B**. Shortest line segments (light blue) between pairs of voxels in top and bottom shells. **C**. Left: Average direction of *N* closest line segments gives local orientation *Ō*. Right: Average length of *N* closest line segments gives local thickness 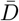. Ratio of distance from top shell *y* to local thickness gives local depth 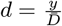. **D**. Illustration of these concepts in the simple geometry of a cube. In addition to top (green) and bottom (pink) shells, sides are labeled (light blue). Shortest line segments are all parallel and have the same orientation *O* and same height *D*. There is a single line segment passing through every voxel center.

Having done this, we proceed to find for each voxel in the top shell, the closest voxel in the bottom shell, and vice-versa. We define all line segments joining pairs of voxel centers (Fig. 1B) and use them to define the local orientation of the principal axis and the local thickness of the source volume (measured along the principal axis) at each voxel (Fig. 1C).

To define the local orientation vector of a voxel, we consider the *N* closest line segments to its center and compute their average direction *Ō*.Conventionally, in order to measure depth, we take the sign of the vector pointing towards the bottom shell. To define the local thickness, we take the average length 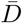 of these *N* closest line segments. Finally, to define the local relative depth *d*, we measure the distance *y* from the voxel center to the top shell along *Ō* and take the quotient with 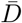, that is 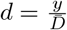.

The number *N* of line segments used per voxel determines to what degree the resulting orientation and depth fields are spatially smoothed. Artifacts in the region annotation atlas and its spatial discretization can lead to sudden jumps in the measurements, making some degree of smoothing necessary; thus, *N* should be chosen depending on voxel resolution and the geometry of the source volume.

### 2.2 Flatmapping

In order to define the other two axes orthogonal to the principal axis in a laminar coordinate system, we introduce a flatmapping algorithm. Intuitively, the process of flatmapping is like squeezing the neocortex by pressing simultaneously from the top and bottom at every point, collapsing all layers into a very thin, but curved sheet of cortex; if we flatten this sheet, and draw coordinate axes on it, we obtain a flatmap. These steps must be done smoothly to preserve *connectedness*, i.e. the 3D subvolume associated with a 2D area has a single component, and *continuity*, i.e. 3D subvolumes associated with neighboring areas are neighbors themselves, in the result.

The input to the flatmapping algorithm is a definition of the principal axis, along which the source volume is to be flattened. Specifically, we require auxiliary atlases of local orientation and relative depth, as are generated by the algorithm in the previous section. The algorithm is flexible, however, and inputs generated by other methods can be used, as long as they provide the same information.

The flatmapping algorithm has two stages (I, II) which can be performed in parallel and converge into a final stage (III). Since the algorithm involves computing depths and orientations at arbitrary locations and not just voxel centers, we define continuous functions through linear interpolation, giving depth *d*(*x*) and orientation *O*(*x*) at any point *x* inside the source volume.

#### 2.2.1 Stage I: Flat mesh generation

##### 1. Definition of projection mesh

The first step is choosing a *projection surface* onto which three-dimensional points are projected. This surface is defined as a level set (isosurface) of the relative depth field, i.e., the set of all points at the same relative depth *d*^∗^. Note that this surface is only defined as an abstraction, as it cannot be computed exactly in a numerical setting.

What can be generated is a *projection mesh*, a triangular mesh that approximates the projection surface. To this end, first we find all voxels (approximately) at a relative depth of *d*^∗^. We do so by defining two binary masks, one with voxels at relative depth less than *d*^∗^ and another with all other voxels, i.e., rest of source volume and outside it, and taking those voxels in the source volume that are at the boundary between the masks.

Then, we apply a surface reconstruction algorithm to generate a triangular mesh from the point cloud of voxel centers. We use the scale-space surface reconstruction algorithm (Lankveld, 2023), as it gives good results and produces a mesh that contains the original points and thus approximately matches the projection surface. Based on the output, manual fixes can be performed to the point cloud to improve the surface reconstruction.

Finally, we increase the number of vertices in the triangular mesh through refinement by subdivision (Geuzaine and Remacle, 2009), that is, introducing new vertices at the midpoints between existing vertices and connecting them to define new triangles. This is needed in order to minimize errors when projecting voxel centers in a following stage. We apply this procedure one or more times to attain a number of vertices in the mesh that is as close as practically possible to the number of voxels in the source volume.

##### 2. Flattening of projection mesh

Flattening of the projection mesh is performed using an area-preserving (authalic) texture mapping algorithm (Saboret et al., 2023). The algorithm takes as input a triangular mesh with one connected component and a continuous boundary, and outputs a triangular mesh in the unit square that is flat and has the same number of points as the input mesh, in a one-to-one correspondence. An area-preserving algorithm is chosen to ensure that the relative sizes of the regions in the source volume is preserved. We call this flattened triangular mesh in [0, 1] *×* [0, 1] the *flat space* and *ℱ* the flattening operator that maps points on the projection mesh to points in flat space.

Since the mesh boundary is mapped to the perimeter of the unit square, there is a choice to be made about which vertices on the mesh boundary map to the corners of the square. We use a simple algorithm that divides the mesh boundary into four parts and uses the breakpoint vertices as corners. In brief, starting at an arbitrary point, we move along the boundary measuring the cumulative length and pick the vertices where quarters of the total length are attained. Given that the choice of corners determines the orientation of the axes in flat space, this step can be optimized depending on the use case, by shifting the starting point along the boundary or even specifying the corners manually.

#### 2.2.2 Stage II: Voxel projection

In this stage we compute a mapping from voxel centers to points on the projection surface. This stage can be performed in parallel to stage I, as it does not require the projection mesh but only the definition of the projection surface (as a level set of the relative depth field).

##### 1. Computation of streamlines

For each voxel *v* in the source volume we generate a *streamline*, a 3D curve that follows the principal axis at every point and connects the top and bottom shells while passing through the voxel center (Fig. 2A, intuitively similar to a radial glial cell). In detail, with the center of *v* as initial position, we compute an integral curve of the orientation field *O* numerically (see details in Supplementary material B). Using a small enough time step, we first iterate in the forward direction until we reach a relative depth of 1 (with *ϵ*_1_ tolerance); then, we iterate in the backward direction until we reach a relative depth of 0 (with *ϵ*_0_ tolerance). Finally, we join these two curve segments and define the streamline associated with voxel *v* as a linear spline *S*_*v*_(*t*), where the parameter *t* ∈ [0, 1] maps monotonically to points along the full curve.

**Figure 2:**
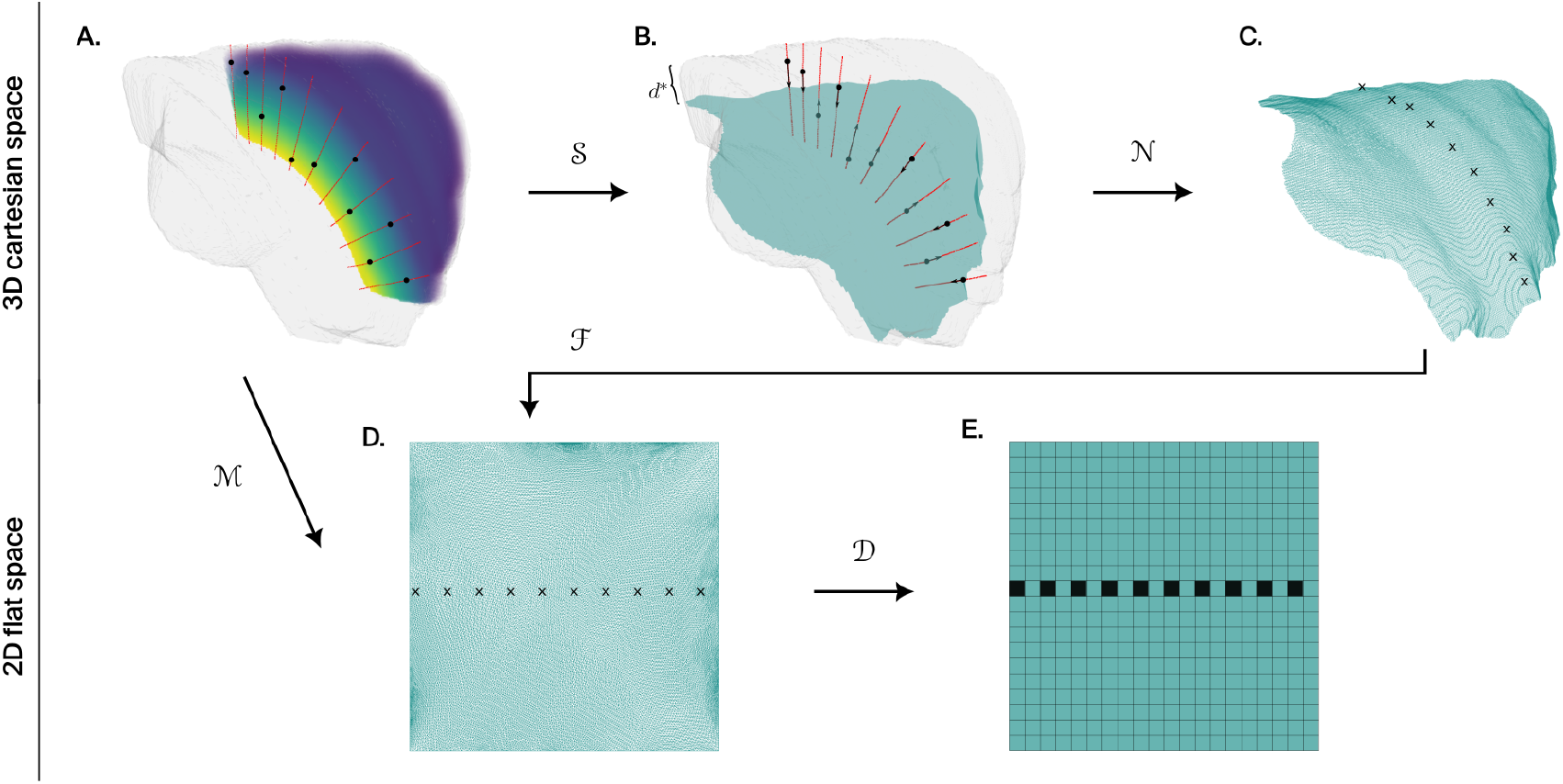
Flatmapping algorithm. **A**. Streamlines (red curves) are computed by numerical integration of the orientation field using voxel centers (black dots) as initial positions. The relative depth field is represented as a colored volume ranging from low depths (purple) to high depths (yellow). **B**. Voxel centers are mapped to the projection surface at relative depth *d*^∗^ (turquoise) following streamlines (black arrows, operator *𝒮*). **C**. Points on the projection surface are mapped to vertices (black crosses) on the reconstructed projection mesh (turquoise wireframe) through nearest-neighbor search (operator *𝒩*). **D**. Authalic (area-preserving) flattening of projection mesh into the unit square (operator *ℱ*). The composite flatmapping operator *ℳ* = *ℱ* (*𝒩* (*𝒮* (·))) maps 3D voxel centers to 2D flat coordinates. **E**. Flatmap discretization by grouping of mesh vertices into pixels of chosen size (operator *𝒟*).

##### 2. Projection of voxel centers

We use the streamline *S*_*v*_(*t*) to find the projection of voxel *v* onto the projection surface (Fig. 2B). Intuitively, we move the voxel center along the streamline until reaching the projection surface. Technically, we define *F* (*t*) = *d*(*S*_*v*_(*t*)) − *d*^∗^ and use a root finding algorithm to find 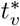, the intersection between the streamline and the projection surface, i.e., the point on the streamline at relative depth *d*^∗^. We call 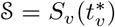 the operator that maps voxel centers to points on the projection surface.

For some voxels, especially at the edges of the source volume, integration of *O* may not reach relative depths of 0 or 1 before leaving the source volume. This may happen due to noise or discontinuities in the orientation field, or truncation of the geometry. These voxels are not *eligible* for flattening and we discard them, since they may not have a projection (if their streamline never attains depth *d*^∗^), and even if they do, the laminar architecture is not preserved along their streamlines.

#### 2.2.3 Stage III: Flatmap generation

Since our ultimate goal is to define a mapping between voxel centers and points in flat space, in the last stage we combine the outputs from the previous stages, namely, the flattened projection mesh and the set of projections of voxel centers onto the projection surface.

##### 1. Mapping to mesh vertices

For each projection of a voxel on the projection surface, we use a spatial nearest-neighbor search algorithm (Tangelder and Fabri, 2023) to find the corresponding closest vertex on the projection mesh. We call this operator *𝒩* (Fig. 2C).

The distance to the nearest neighbor represents the approximation error of the projection mesh to the projection surface at that location. We try to keep these errors low by reconstructing the projection mesh from points near *d*^∗^ and using iterative refinement to increase the number of vertices in the mesh (see stage I.1). However, if these errors are consistently large (half the voxel size or more), this may be indicative of a problem in the definition of the projection mesh.

##### 2. Flatmap generation and discretization

Finally, thanks to the one-to-one correspondence between vertices in the projection mesh and flat space, we map each eligible voxel to its *flat projection*. Mathematically, this is represented by the composite *flatmap* operator *ℳ* = *ℱ*°*𝒩* °*𝒮*. In summary, each voxel center *v* is moved following its streamline to the projection surface, then to the nearest vertex on the projection mesh, and finally to the corresponding vertex in flat space.

The coordinates of the flat projection are called the *flat coordinates* of a voxel. The highest resolution of a flatmap is determined by the number of vertices in the projection mesh, and we call this the *native resolution*. For certain applications, however, it is desirable to have a regular discretization of the flat space at a lower *pixel resolution N*, meaning that floating-point coordinates in [0, 1] are binned into integer indices from 0 to *N* − 1 in both axes (Fig. 2E).

In general, many voxels will have the same flat projection. The set of voxels that map to a given point in flat space, whether at native or pixel resolution, is called the *pre-image* of that point, and can be represented by the inverse operator *ℳ*^−1^. Pixels with empty pre-images, i.e. that no voxels map to, may be undesirable depending on the application and can be avoided by a proper choice of pixel resolution.

### 2.3 Distance and area measurements in flat space

Given that the flattening algorithm always maps the projection mesh to the unit square, flat coordinates are normalized to [0, 1] and distance units are lost. In order to recover distance units in flat space, we compute pre-image radius at a given pixel resolution (see 2.5.3) and then establish a linear fit between mean pre-image radius (in *µ*m) and pixel size. The slope of the linear fit can be used as a proportionality factor between normalized distances in flat space and real distances in Cartesian space. This definition assumes isotropy in flat space, which may or may not be achieved depending on the aspect ratio of the projection mesh.

To measure areas in flat space, since the flattening algorithm is area-preserving, it suffices to use the original area of the projection mesh as a proportionality factor.

### 2.4 Characterization of depth and orientation fields

In order to quantify their smoothness, we compute the Laplacian of the depth field and the divergence of the orientation field at every voxel, and check how close these quantities are to zero. That is, we measure how close the depth and orientation fields are to being a homogeneous solution to the heat equation and its gradient, respectively. We compute statistics of these quantities over all interior voxels (not in the top, bottom or sides), to reduce bias due to discrete numerical derivative artifacts at the boundaries of the source volume.

### 2.5 Flatmap characterization

As mentioned before, important properties of a flatmap are: continuity, connectedness, and mapping of 2D areas to 3D subvolumes with preserved layer structure. We define some metrics to characterize how close the output of our flatmapping algorithm is to having these desired properties. Some are global or per-voxel metrics, but otherwise they are defined for each pixel at a given pixel resolution. These per-pixel metrics characterize the structure of the pre-images and can be further analyzed in terms of statistics computed over all pixels and across pixel resolutions.

#### 2.5.1 Global metrics

##### Coverage

Fraction of eligible voxels in the source volume, i.e., number of voxels having a flat projection over total number of voxels. Under normal circumstances this value is expected to be close to 1, with some non-eligible voxels at the edges of the source volume. A value much lower than 1 may be indicative of issues with the orientation field or the integration of streamlines.

##### Usage fraction

Fraction of pixels in flat space with a non-empty pre-image, i.e., with at least one voxel mapping to them, computed at some pixel resolution. At low resolution the usage fraction is almost certainly equal to 1, whereas at higher resolutions holes eventually appear and the usage fraction becomes lower than 1. The optimal pixel resolution depends on the application, and this metric can help in choosing it. At native resolution, the usage fraction is the ratio of the number of vertices with non-empty pre-images to the total number of vertices in the mesh.

#### 2.5.2 Per-voxel metrics

##### Orthogonality

Degree to which the principal axis is orthogonal to the flatmap axes at each voxel. We compute this by first taking the gradient of both flat coordinates considered as scalar fields in Cartesian space. Then, we compute the cross product of these two gradients and choose the sense towards the bottom shell. Finally, we measure the cosine of the angle between this vector and the orientation vector by taking the dot product after normalization. A value of 1 indicates that the principal axis is locally orthogonal to the flatmap, whereas a lower value indicates some degree of non-orthogonality. If this quantity is negative it means the cross product points in the opposite direction as the orientation vector, and a sign flip is needed for proper comparison.

#### 2.5.3 Per-pixel metrics

##### Pre-image size

Number of eligible voxels in the pre-image of a pixel (Fig. 3A).

**Figure 3:**
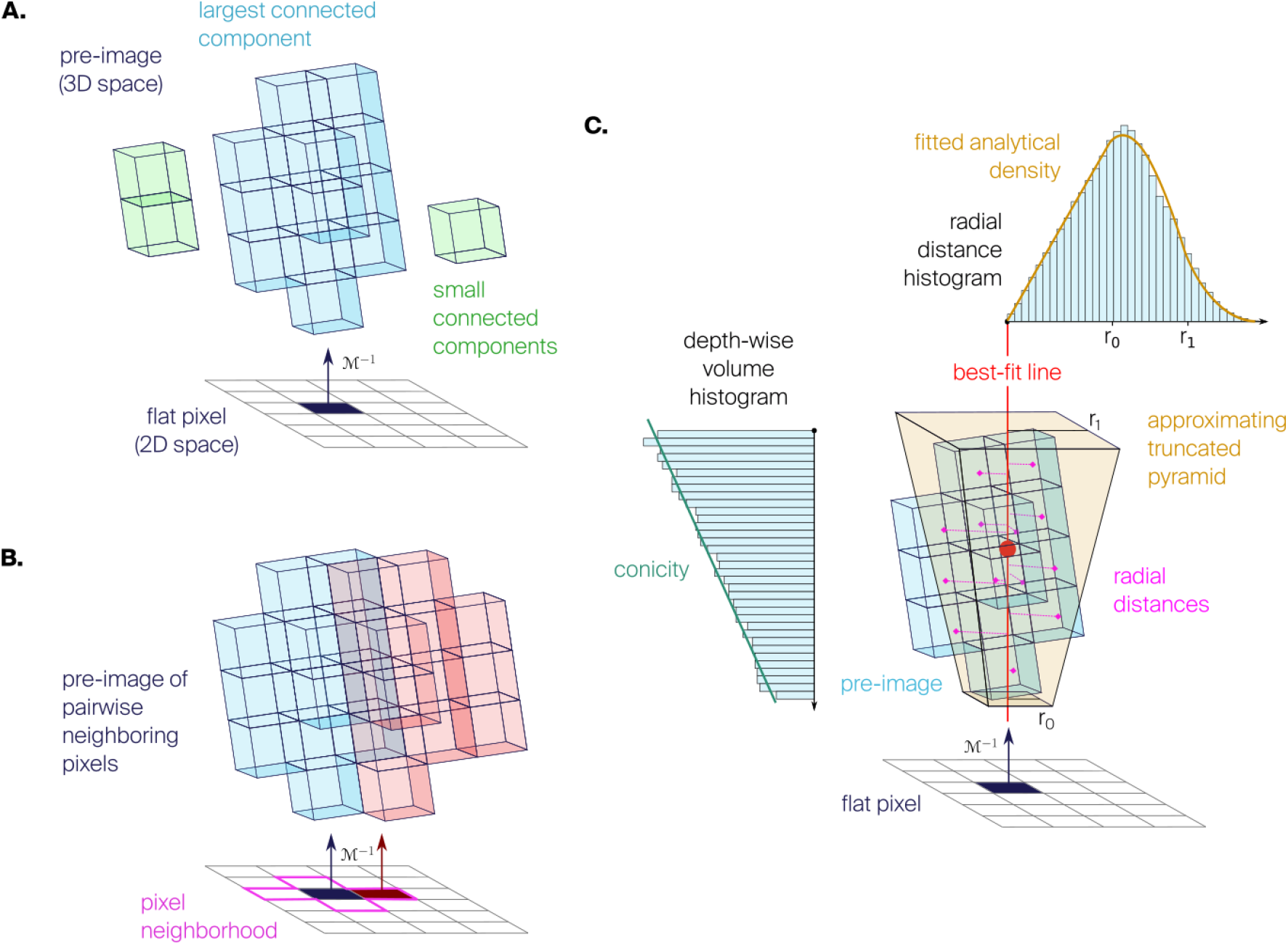
Flatmap characterization. **A**. Pre-image size and connectedness. The pre-image may consist of one (blue) or more (green) connected components. **B**. Pre-image continuity for a pair of neighboring pixels (red, blue). The metric is evaluated over the 4 pairs in the pixel’s neighborhood (magenta). **C**. Pre-image radius is computed using the radii of the best approximating truncated square pyramid, determined by fitting the analytical density to the histogram of radial distances from the best-fit line. Pre-image conicity is the slope of a linear fit to the volume histogram as a function of depth.

##### Pre-image size uniformity

Percentage of eligible voxels in the pre-image of a pixel, relative to the number expected from pixel area alone if voxels were distributed uniformly throughout. In the ideal case, this metric would reflect the purely geometrical variations in thickness of the source volume, as this would determine how many voxels project to a pixel while following streamlines locally. However, noise and artifacts in the orientation field can lead to off-site streamlines that artificially increase the pre-image size at locations far from the neighborhood of a voxel, leading to artifacts visible as non-uniformities in pre-image size.

##### Pre-image connectedness

Ratio of the number of voxels in the largest connected component of the pre-image of a pixel, to the number of voxels in the entire pre-image (Fig. 3A). We consider two voxels to be connected if one is inside the 3×3×3 neighborhood centered on the other (26 adjacent voxels). This metric quantifies how much the pre-image consists of a single chunk vs multiple separate pieces. By taking the ratio of the largest component to the total, instead of the number of connected components, we avoid bias due to possibly one large and many small components (even single voxels). At native or high pixel resolutions, pre-images are expected to be disconnected since the noise in the geometry and the orientation field are more evident than at lower resolutions.

##### Pairwise pre-image continuity

Same as pre-image connectedness, but computed for the combined pre-image of a pair of adjacent pixels. Given there are four adjacent pixels in a pixel’s neighborhood, we take the minimum value (worst case) over all four pairs (Fig. 3B). This metric provides a measure of local continuity in the sense that moving from one pixel to the next in flat space should represent a corresponding movement between contiguous places in Cartesian space. This is important, as otherwise it would not be possible to relate close points in Cartesian space to close points in flat space. A discontinuous flatmap would be indicative of issues with the orientation field, as streamlines should vary smoothly and not cross or deviate from their neighbors.

##### Pre-image conicity

Slope of a line fitted to the depth-wise volume histogram of the pre-image of a pixel (Fig. 3C). When conicity is non-zero, the pre-image has relatively more or less volume at higher depths (for positive/negative values, respectively), and thus resembles a cone. When conicity is zero, the pre-image has a uniform volume distribution along depth, resembling a cylinder.

##### Pre-image radius

Extent of the pre-image of a pixel in the directions orthogonal to its main axis. To give some intuition, if the pre-image was a cylinder, this would be its radius. However, for a square pixel the pre-image looks more like a truncated square pyramid (due to conicity), so we require a way to measure the extent of such an object. We approximate it in three steps (Fig. 3C). First, the center of the pre-image is calculated as the centroid of its voxels. The main axis of the pre-image is defined as a line passing through its center and having the direction that minimizes the sum of square distances to all of its voxels, i.e., the first principal component of the voxels’ coordinates. Then, the radial distance from the main axis to each voxel *r*(*υ*) is calculated, and its histogram is fitted using the analytical formula for a truncated square pyramid of radii *r*_0_ and *r*_1_ (see Supplementary material D). Finally, the pre-image radius is defined as the mean of the fitted radii (*r*_0_ + *r*_1_)*/*2.

### 2.6 Applications of flatmaps

#### 2.6.1 Volume decomposition

The flatmap can be used to decompose the source volume into uniform and non-overlapping subvolumes that preserve the laminar structure, e.g., each sampling the complete stack of cortical layers in the case of the neocortex.

In general, to define a subvolume we first draw a shape in flat space (preferably at native resolution) and take all points that are inside this shape. Then, we take the pre-images of all these points and consider their union, i.e., the set of all voxels that under the flatmap operator project to this flat shape. The resulting set of voxels describes a 3D volume that “follows” the streamlines and keeps the laminar structure intact.

Now, if we draw a tiling in flat space (e.g., square or hexagonal grid) and consider the subvolumes corresponding to all elements of the tiling, we obtain a decomposition of the entire source volume. These subvolumes are non-overlapping as long as the flat shapes are non-overlapping, as is the case for a plane tiling, and their properties can be characterized using the metrics described above (taking “per-pixel” to mean “per-element of the tiling”).

#### 2.6.2 Flat views of three-dimensional data

Any quantity defined for the voxels in the source volume, such as region annotations or injection densities, can be assigned to their flat projections, resulting in a *flat view* (2D representation of 3D data) of such a quantity. Because of this decrease in dimensionality, some aggregation function must be applied to assign the value at a flat point using the combined values of the voxels in its pre-image. Once we have the aggregated values, we can use a color palette to obtain a heatmap of the quantity of interest.

We colloquially call *flatmap* to the flat view of region annotations, and we use the mode (most common value) as the aggregation function. For flat views of other volumetric datasets, we typically use the mean or maximum (similar to a maximum intensity projection) as aggregation function.

#### 2.6.3 Annotation of barrel columns in the mouse isocortex

A variant of the volume decomposition procedure can be used to generate annotations of barrel columns in the mouse isocortex. For this use case, the shapes drawn in flat space are the outlines of the individual barrels, generated as follows.

Firstly, barrel annotations were extracted based on the “average template” two-photon tomography dataset part of CCFv3 (Wang et al., 2020), where the anatomical features of the barrel and septum separation are visually apparent.

Secondly, we performed a semi-automatic segmentation in ITK-SNAP using the Active Contour Snake algorithm (Yushkevich, Y. Gao, and Gerig, 2016), and obtained volumes for 33 ellipsoidal barrels in the left hemisphere. We annotated the 33 barrels as separate structures with labels of their corresponding rows and arcs. Segmentation of additional barrels was unsuccessful due to the limited signal-to-noise ratio of the average template dataset.

Thirdly, the new annotations were projected onto flat space using the flatmap of the mouse isocortex (Fig. 6A). We used the flat positions to further refine the annotations by reassigning mislabeled voxels to the label of their nearest neighbor.

Lastly, using the final annotations, we computed the convex hull of all flat positions belonging to a barrel and took the pre-image of all points inside it. This way we obtained cylinder-shaped barrels and barrel columns. The originally ellipsoidal barrels are now extended to span the complete height of layer 4 (Fig. 6B).

#### 2.6.4 Flat views of long-range axons in the mouse isocortex

Any morphology reconstruction with points inside the source volume can be visualized in flat space. This is done by simply taking the flat projection of the points representing the soma and neurites, and plotting them over the flatmap.

In particular, for visualization of long-range axons in the mouse isocortex, we downloaded in bulk the latest MouseLight data release from ml-neuronbrowser.janelia.org (dated 2022.02.28) and worked with JSON morphologies in the CCFv3 space. We then plotted the flat projections of the soma and axon points on top of our flatmap of mouse isocortex for both hemispheres (see Fig. 7).

## 3 Results

### 3.1 Enhancement of rat somatosensory cortex atlas

We applied our methods to an atlas of rat somatosensory cortex with 10 annotated regions (Fig. 4A), based on a well-known printed atlas of the rat brain (Paxinos and Watson, 2007). Details of the creation of this atlas and a description of the region labels can be found in Supplementary material A.

**Figure 4:**
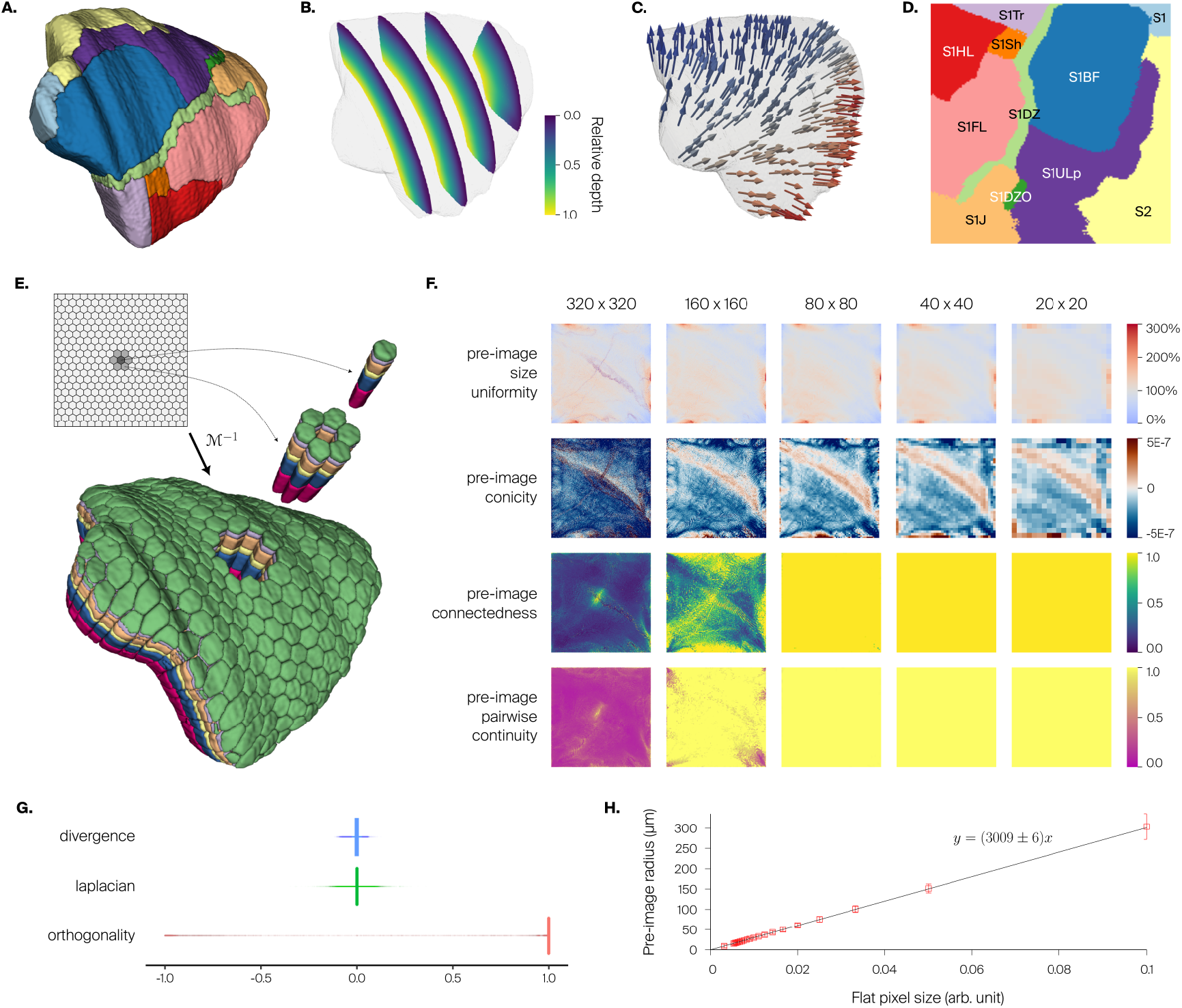
Enhancement of rat somatosensory cortex atlas. **A**. 3D atlas with region annotations for one hemisphere. **B**. Relative depth field, represented as sections with colormap giving relative cortical depth. **C**. Orientation field, represented as small 3D arrows (sample). **D**. Flatmap of somatosensory regions in one hemisphere (same colors as in A), with labels. **E**. Decomposition of atlas volume into hexagonal subvolumes using the inverse flatmap operator *ℳ*^−1^. Preservation of layer structure (in colors, from L1 in green to L6 in magenta) is evident in hexagonal columns shown separated from the whole. Inset shows hexagonal grid drawn on flat space, with highlight of central hexagon (dark gray) and surrounding hexagons (light gray). **F**. Quality metrics as a function of pixel resolution. For pre-image size uniformity, the color bar indicates the proportion of voxels mapped to a pixel compared to that expected from pixel area alone in the perfectly uniform case; values at or above 300% are colored the same. We show power-of-two pixel resolutions to have nice sub-sampling in plots. In general, degradation is visible near the edges. **G**. Distributions of values of per-voxel metrics: divergence of orientation field, Laplacian of relative depth field, and orthogonality between principal axis and flatmap axes. **H**. Distance metric obtained from linear fit of flat pixel size vs. pre-image radius (mean and std. dev.).

After manual expert selection of the top and bottom shells, we generated a relative depth field (Fig. 4B) and orientation field (Fig. 4C) sampling *N* = 1000 nearest line segments at each voxel.

For flatmapping, the source volume consists of all somatosensory regions, encompassing 1, 314, 885 voxels. The projection surface is taken as the isosurface of the relative depth field at *d*^∗^ = 0.5. The reconstructed projection mesh has 23, 314 vertices and an area of 40.606 mm^2^, and it was refined three successive times into 1, 492, 096 vertices. The resulting flatmap (Fig. 4D) shows clearly delineated regions. It has a usage fraction at native resolution of 0.581 with a mean pre-image size of 1.5 voxels, and the coverage is 0.988 with only 15, 677 non-eligible voxels.

We show a decomposition of the source volume into hexagonal subvolumes (see Methods 2.6.1; Fig. 4E). Note how the hexagonal grid defined in flat space is well transferred to Cartesian space. Furthermore, the subvolumes are non-overlapping and have an intact layer structure, resembling cortical columns.

We investigated the impact of pixel resolution on the properties of pre-images by computing a number of metrics (see Methods 2.5; Fig. 4F). We show resolutions from 320 × 320 pixels, which is closer to native resolution and is better for drawing on flat space, to 20 × 20 pixels, which is close to the resolution at which we decompose the source volume, e.g., into hexagonal columns as above. We found the resulting pre-image sizes to be relatively uniform at all resolutions, indicating that different parts of the flat space are comparable in how they sample the Cartesian space. However, at a resolution of 320 × 320 pixels, the remaining variability still resulted in a number of empty pre-images, i.e. holes. Pre-image conicity was noisier at high resolutions, probably resulting from the uneven surface structure of the source volume (Fig. 4A vs. 4F, third row). At resolutions below 80 × 80 pixels, this property was smoother and more uniform. Pre-image connectedness and continuity were largely ensured at resolutions of 80 × 80 or lower, but fell apart for higher resolutions. For all metrics, the locations of largest non-uniformity were at the edges of the geometry.

Per-voxel metrics show good results (Fig. 4G). Computing the Laplacian of the relative depth field as a measure of smoothness shows good results with a very narrow distribution symmetrical around zero and median value of 0.0003 (0.003 inter-quartile range (IQR)). This value is small compared to the range [0, 1] of the relative depth field. Computing the divergence of the orientation field as a measure of smoothness also shows good results with a slightly wide distribution symmetrical around zero and median value of 0.0 (0.013 IQR). We also compute orthogonality of the local orientation vector to the local flatmap axes, obtaining good results with a very sharp distribution near one and median value of 0.999 (0.001 IQR). This demonstrates that the decomposition into one principal axis and two axes orthogonal to it was successful.

Finally, we obtained a linear fit of flat pixel size vs. pre-image radius (Fig. 4H), leading to a distance metric in flat space of 3009 *±* 6 *µ*m per flat coordinate unit.

### 3.2 Enhancement of mouse isocortex atlas

Owing to the flexibility of our flatmapping method, we use existing datasets from the Allen Institute Mouse Brain Atlas to generate a flatmap of the entire mouse isocortex. We take the solution to Laplace’s equation available in the CCFv3 2017 release at 10 um resolution (`laplacian_10.nrrd`), and use it as relative depth field (Fig. 5B). Then, we compute its gradient numerically and use it as orientation field (Fig. 5C). Finally, we apply the same algorithm as before, except we use an iterative version of the authalic mapping algorithm (Saboret et al., 2023).

**Figure 5:**
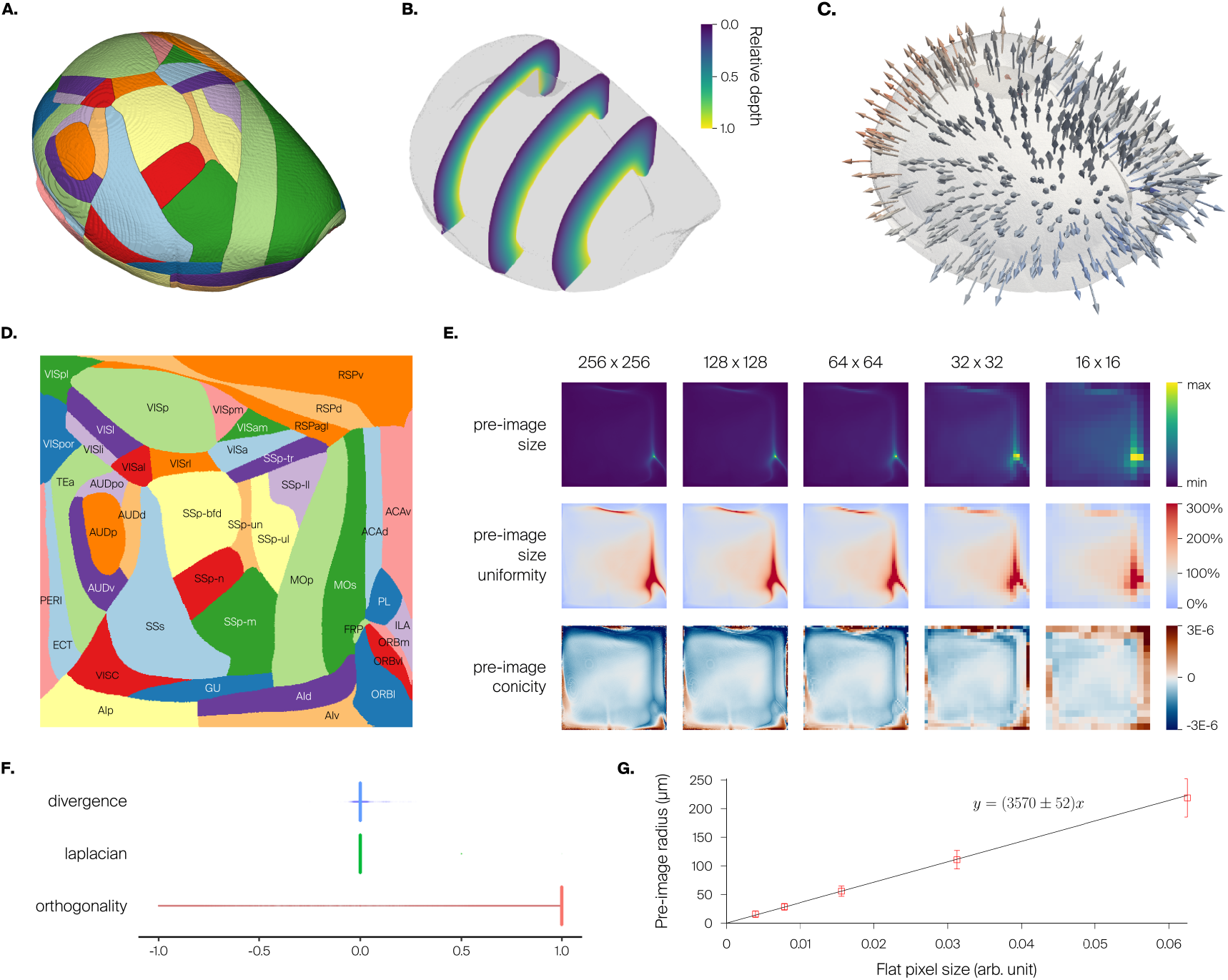
Enhancement of mouse isocortex atlas. **A**. 3D atlas with region annotations for one hemisphere. **B**. Solution to Laplace’s equation as depth field, represented as slices with colormap giving relative cortical depth. **C**. Gradient of the solution to Laplace equation as orientation field, represented as small 3D arrows. **D**. Flatmap of all isocortex regions in one hemisphere (same colors as in A), with labels. **E**. Quality metrics as a function of pixel resolution. For pre-image size, the color bar indicates the range of values present at each pixel resolution, and cannot be quantitatively compared across resolutions. For pre-image size uniformity, the color bar indicates the proportion of voxels mapped to a pixel compared to that expected from the pixel area alone in the perfectly uniform case; values at or above 300% are colored the same. We show power-of-two resolutions to have nice sub-sampling in plots. “Hotspots” in pre-image size are visible in the frontal pole and other regions of high cortical curvature. **F**. Distributions of values of per-voxel metrics: divergence of orientation field, Laplacian of relative depth field, and orthogonality between the principal axis and flatmap axes. **G**. Distance metric obtained from a linear fit of flat pixel size vs. pre-image radius.

We took all 43 regions under “Isocortex” in the hierarchy of the CCFv3 as the source volume (Fig. 5A), encompassing 61, 945, 750 voxels. The availability of top, bottom and side shells in the CCFv3 release would allow us to derive the relative depth and orientation fields with our methods. However, we prefer to use the already available solution to the Laplace equation to showcase integration with external datasets derived using different methods from ours. When computing the gradient, we extend the solution one voxel towards the outside to reduce edge artifacts in the numerical derivatives. We take as projection surface the isosurface of the Laplacian solution at *d*^∗^ = 0.5. The reconstructed projection mesh has 437, 186 vertices and an area of 51.652 mm^2^, and it was refined three successive times into 27, 942, 965 vertices.

The resulting flatmap shows clearly delineated regions (Fig. 5D). It has a usage fraction at native resolution of 0.833 with a mean pre-image size of 2.7 voxels, and the coverage is 0.997 with only 213, 025 non-eligible voxels.

In terms of pre-image properties (Fig. 5E), we found that the resulting flatmap was almost completely connected and continuous without holes at resolutions up to 256 × 256 pixels (not shown for simplicity). Higher pixel resolutions could not be investigated due to computational costs stemming from the high resolution (10*µm*) of the input atlas, but based on the behavior of the metrics for the rat flatmap above, a breakdown of connectedness would be expected at a resolution of 1024 × 1024 pixels, and the appearance of holes would be expected at a resolution of 2048 × 2048 pixels. Pre-image conicity was smoother than for rat at high resolutions, reflecting the smoothness of the input atlas. One noteworthy feature visible in the pre-image size and uniformity at all pixel resolutions is the presence of “hotspots” in the frontal pole region and in regions along the edge where the cortical hemispheres meet (compare to Fig. S2B in Wang et al., 2020). These are places of high curvature in the cortex, where due to convergence of orientation vectors many voxels are mapped to the same pixel.

The per-voxel metrics showed excellent results (Fig. 5F). As expected, the Laplacian of the depth field (itself a solution to Laplace equation) was very close to zero, with a median of 1.0 × 10^−5^ (2 × 10^−5^ IQR), and the divergence of the orientation field was correspondingly also very close to zero, with a median of 1.1 × 10^−5^ (2 × 10^−5^ IQR). Orthogonality showed once more a successful decomposition into a principal axis and two orthogonal axes, with a median of 0.999 (0.0013 IQR).

Finally, we obtained a distance metric in flat space with 3570 *±* 52 *µ*m per flat coordinate unit (Fig. 5G).

Since the CCFv3 atlas is perfectly symmetrical with respect to the midline, we extended our flatmap to both hemispheres by mirroring, adjusting the flat coordinates to have *x* ranging in [0, 1] for the right hemisphere and [1, 2] for the left hemisphere. We use this two-hemisphere flatmap for data visualization (Fig. 7C, E).

**Figure 6:**
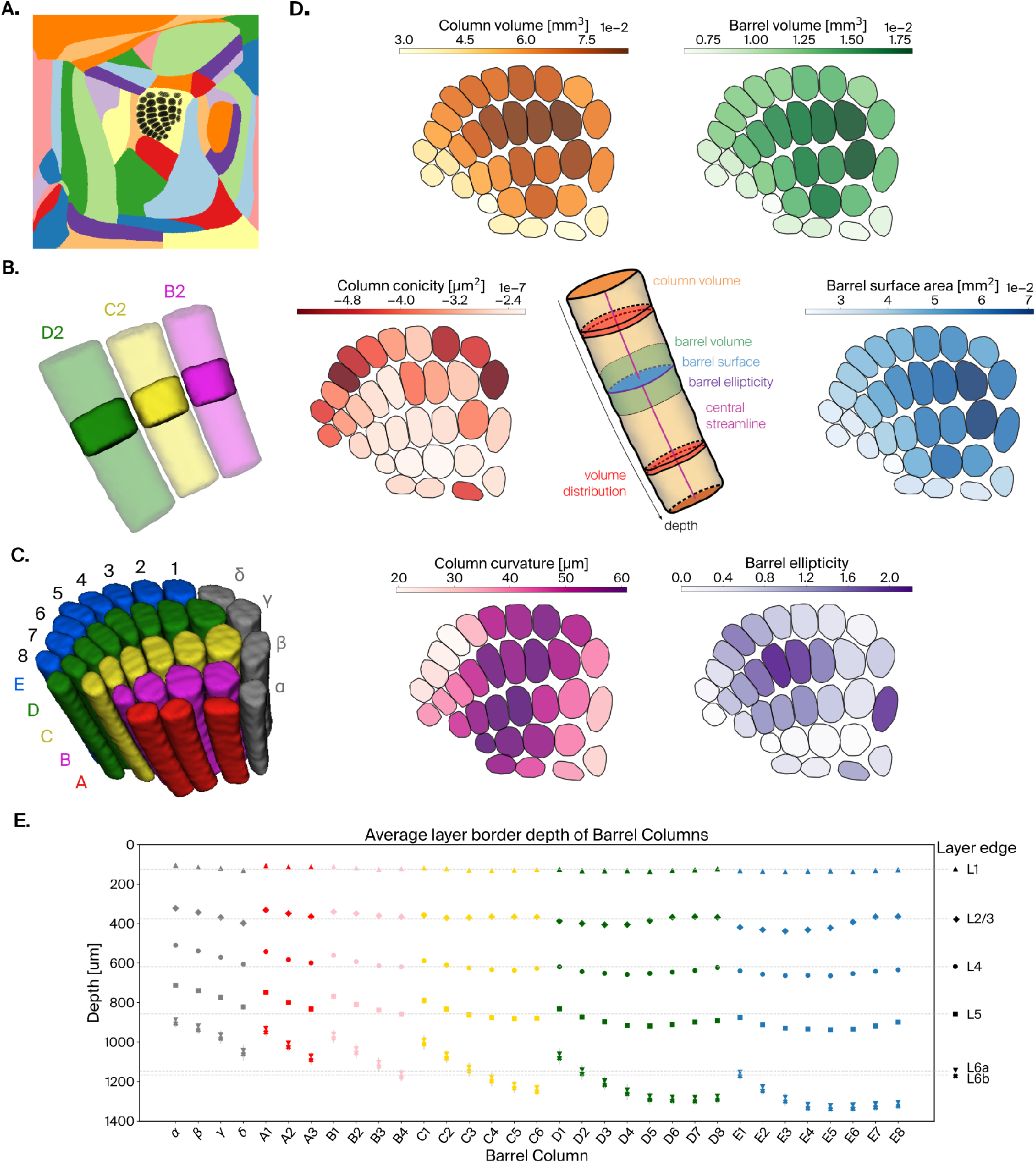
Enhancement of mouse barrel cortex atlas. **A**. Flat projections of annotated barrel voxels (see Methods, black) in SSp-bfd, shown in the context of the full mouse isocortex. **B**. Barrel volumes of 2-row (solid) and their corresponding columns, defined as the pre-images of their flat projections (semi-transparent). **C**. Extracted barrel columns for all annotated barrels, barrel rows represented by the same color. **D**. Barrel and barrel columns metrics. The center schematic illustrates how they are defined. Followed by heatmaps of their values plotted on flat projections of barrels. The metrics are (clockwise): *Barrel volume*, shown as an outline (green) in the schematic. *Barrel surface area*, shown as a cross-section (blue) in the schematic. *Barrel ellipticity*, computed as the ratio of major to minor axis of en ellipse fitted to the flat projection of a barrel, represented by the perimeter (purple) in the schematic. *Column volume*, shown as a volume (orange) in the schematic. *Column curvature*, defined as maximal displacement from a straight line fitted to the barrel streamline, shown as a line (pink) in the schematic. *Column conicity*, computed as the slope of a linear fit to the depth-wise volume histogram, represented by two volume bins (red) in the schematic. **E**. Average layer border depth for all layers in each of the barrel columns. Grey lines indicated mean layer depths across all barrel columns.

**Figure 7:**
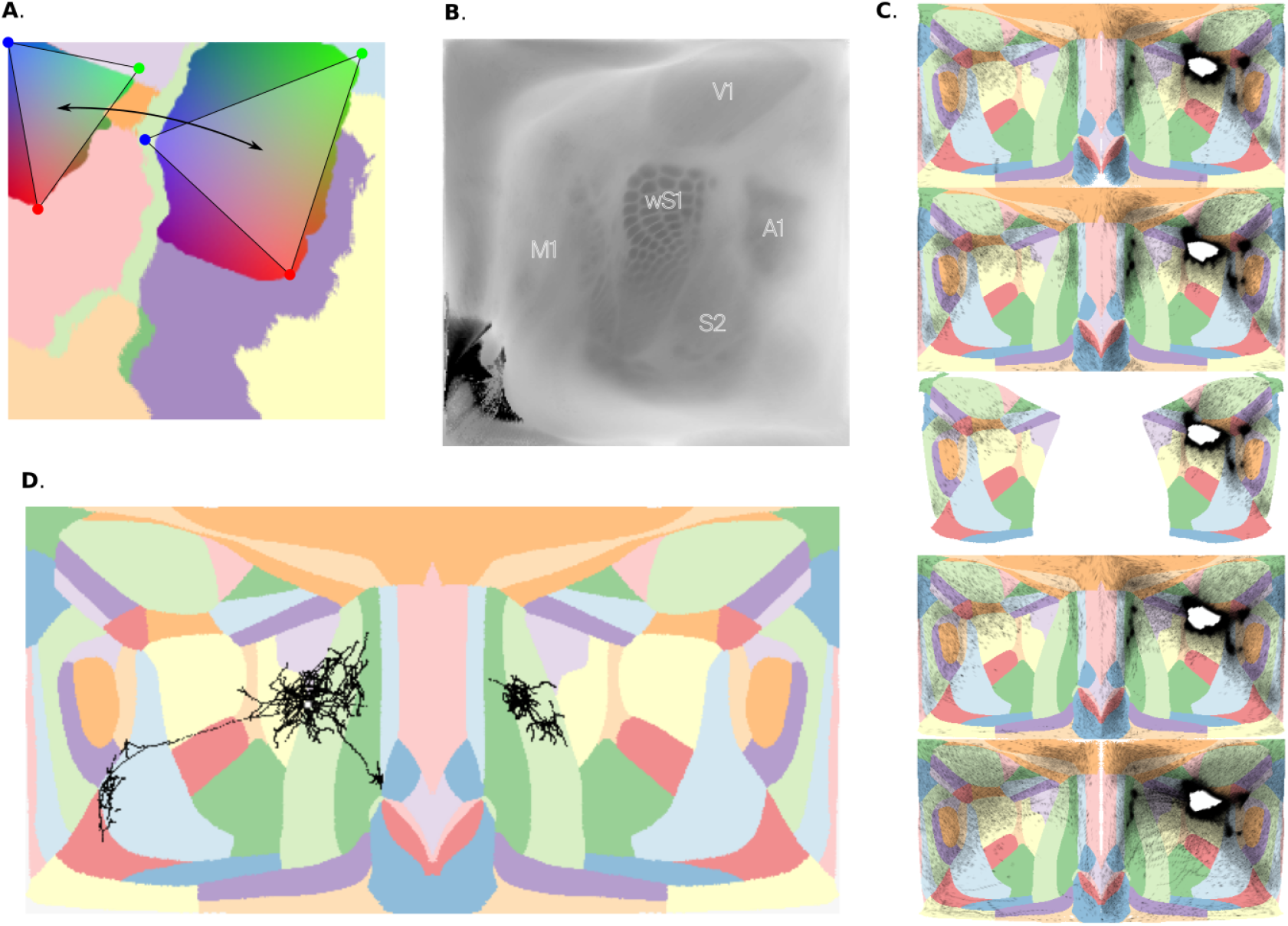
Applications of flatmaps. **A**. Topographical mapping between two rat somatosensory regions. Colors represent barycentric coordinates of triangles contained in each region (red, green and blue vertices). Locations with the same color connect to each other. **B**. Flat view of the CCFv3 average template of mouse isocortex. **C**. Flat views of injection (white) and projection density (black) across cortical layers in both hemispheres of mouse isocortex. Per-layer maps are stacked, and correspond to L1, L2/3, L4, L5 and L6. **D**. Flat view of long-range axons (black points) of a neuron with soma (white dot) in primary motor cortex (MOp) of the left hemisphere, near the border with lower limb primary somatosensory cortex (SSp-ll).

### 3.3 Enhancement of mouse barrel cortex atlas

We segmented well-defined barrels from the average template dataset in CCFv3 (Wang et al., 2020) and produced an annotation of the posterior medial barrel field, with 33 individually labelled barrels (Welker and Woolsey, 1974; Simons and Woolsey, 1979). We then used an approach based on flat projection of barrel annotations (Fig. 6A) and taking their pre-images (see Methods 2.6.3), to produce annotations of barrel columns in the reference coordinate space of the Allen CCFv3 mouse brain atlas (at 10 *µ*m resolution).

Our efforts allowed us to derive annotations with the following properties: barrels are cylindrical, span the full height of layer 4 and do not overlap, and barrel columns follow the curvature of the cortex. Thanks to these properties, where neighboring barrels are separated by the septum, the associated barrel columns are also separated; where barrels contact each other, their associated barrel columns are also in contact, but never overlap throughout the entire depth (Fig. 6B and C).

To further characterize the geometry of the new annotations, we computed a number of anatomical metrics (Fig. 6D). For the surface of each barrel we considered the convex hull of the flat projections of the barrel voxels and calculated their area as previously described (see Methods 2.3). To characterize the shape of the barrels we measure their ellipticity, computed as the ratio of major to minor axis of a confidence ellipsoid fitted to the flat projection of a barrel. The resulting surfaces have an elongated shape, in some cases twice as wide along an arc than along a row, in line with (Welker and Woolsey, 1974). Furthermore, barrels within a row are very close to each other, often touch, but do not overlap, whilst big septal areas can be observed between the rows.

We then calculated the volumes of barrels and barrel columns, based on the voxel count and voxel size. In terms of volume and surface area barrels were generally smaller towards the periphery, with barrel D1 being the largest. Column volume was strongly, but not completely dependent on barrel surface area. An additional factor was the increase of cortical thickness from row A to row E.

Additionally, we characterized how much the columns deviate from right cylinders, measuring their *conicity* and *curvature*. The former is defined as the slope of a linear fit to the depth-wise volume histogram (see Methods 2.5.3). The latter metric is defined as the maximal displacement from a straight line fitted to the barrel streamline, at the top or bottom of the barrel column.

Intuitively we measure how much the end of the barrel column’s streamline deviates from a straight line. For example, from layer 4 the C2 barrel curves 50 *µm* towards the C1 barrel column compared to a corresponding representation as a right cylinder. As a result of the global curvature of the cortical volume, columns were wider at the top than at the bottom. The strength of this effect was independent of column curvature. The approximated radius of the C2 barrel column narrowed by 32 *µ*m over the entire depth.

Finally, we found the layer borders and quantified them depth-wise along the barrel columns (Fig. 6E). We found that the barrel columns increase in height from row A to E. We can also observe a pattern of height increasing with the number of the arc. However, the relative layer depths are largely preserved across the barrel columns.

### 3.4 Applications of flatmaps

The flatmaps obtained in the previous section have multiple applications, a few of which we highlight here.

#### 3.4.1 Topographical mapping of long-range connectivity

A flat view of region annotations can be used to visualize and describe the topographical mapping of connections between cortical areas. While the spatial targeting of cortico-cortical connections is often expressed in terms of laminar synapse profiles (i.e., specificity along an axis orthogonal to layer boundaries), axons from individual neurons also target specific locations along the other two axes, and this can be expressed as a topographical mapping between points in flat space. In our flatmap of rat somatosensory cortex, we parameterized the topographic mapping by using triangles defining local barycentric coordinates per region, and then considering pairs of triangles so that neurons at a given location in one triangle predominately project to the corresponding point on the other triangle (Fig. 7A).

#### 3.4.2 Volumetric data visualization

A flatmap allows us to visualize volumetric data in 2D, as long as the data is registered to the same brain atlas that was flatmapped. We provide two examples obtained using our flatmap of mouse isocortex.

We show a flat view (Fig. 7B) of the average template of the adult mouse brain in CCFv3 (Wang et al., 2020). This dataset was constructed by interpolation of tissue auto-fluorescence data from high-resolution serial two-photon tomography images of 1,675 young adult mouse brains. The flat view was obtained using the maximum as aggregation function, resulting in a flat maximum intensity projection (fMIP). Indeed, the resulting image is very similar to a cytochrome oxidase staining of mechanically flattened cortex (compare Fig. 7B to Fig. 1 of Airey et al., 2005).

We also show connectivity data for a single experiment (ID 126907302) from the Allen Mouse Connectivity Atlas (Oh et al., 2014). Specifically, we show the injection and projection densities of connections originating mainly from the SSp-bfd region, but with slight injection density in the rostrolateral visual area (VISrl). In order to visualize information with laminar specificity, we split the flat view by layer (Fig. 7C). In the images we can see the strongest cortical targets were parts of the ipsilateral secondary somatosensory and motor areas, the dorsal auditory area (AUDd), and the neighboring anterolateral (VISal) and anterior (VISa) visual areas. Weaker targets, which suffer from high noise, were the remaining visual and auditory regions, and some retrosplenial, temporal and orbital areas. Other prefrontal and anteromedial areas, including most of primary somatosensory cortex, were largely avoided. Layer-wise, we observe that both injection and projection densities spread to all layers, but with slightly higher projection density in infragranular layers.

#### 3.4.3 Long-range axon visualization

We used our flatmap of mouse isocortex to visualize the anatomy and targeting of long-range axons, using data from Janelia MouseLight (mouselight.janelia.org; Economo et al., 2016). Specifically, we show an exemplary mouse axon (ID AA0002) from a cell with soma in the primary motor cortex (MOp) near the border with primary somatosensory lower limb region (SSp-ll). Our approach depicts the targeting and branching structure of all axon collaterals in a single plot. Furthermore, it visualizes the horizontal targeting of axons independently from the vertical one, i.e. independently from their laminar termination profiles. In this flat view (Fig. 7D) we can observe branches innervating nearby parts of SSp-ll, as well as ipsilateral secondary somatosensory and motor regions with high specificity. A single branch extends ipsilaterally to secondary somatosensory cortex while other branches innervate motor regions in the contralateral hemisphere. As the flatmap covers only the isocortex, parts of the axon leaving this part of the brain are not visible. Therefore the contralateral innervating parts appear separated from the rest and branches innervating subcortical structures are not represented at all.

## 4 Discussion

We have demonstrated a method to enhance brain atlases with three-dimensional coordinate systems adapted to the geometry of layered brain regions. These *laminar coordinate systems* consist of auxiliary atlases describing the principal axis (local relative depth and orientation) and flatmap (other two horizontal axes) of the region of interest. Furthermore, we have defined a set of metrics to characterize the quality of flatmaps, and introduced several applications highlighting the utility of laminar coordinate systems for data visualization and data-driven modeling.

In particular, we applied our method to an atlas of rat somatosensory cortex based on Paxinos and Watson, 2007 (see Supplementary material A), enhancing it with a laminar coordinate system adapted to the geometry of this region. We also applied our method to a well-known atlas of the mouse brain (Wang et al., 2020), enhancing it with a flatmap of the whole isocortex, which we used to produce new annotations of 33 individual barrels and barrel columns in the barrel cortex.

Importantly, we have made an implementation of all our methods publicly available as free software, as well as the enhanced atlases and flatmaps obtained in our results (see Code and data availability).

While this is not the first time the concept of a flatmap has been introduced, we have described a general method that applies to any brain region with a principal axis of organization. Unlike flatmaps of a more schematic nature (Hahn, Swanson, et al., 2021), the type of data-driven flatmap produced by our method is adapted to the geometry of the voxelized atlas it is derived from. This enables a direct representation of data registered to the atlas (see 3.4), similar to the flatmap developed in Wang et al., 2020 for mouse isocortex. Our approach differs from theirs as we use an area-preserving algorithm, instead of distance-preserving, to parameterize the flat space. Our methodology also diverges from that of Diedrichsen and Zotow, 2015 as our flat space comprises a continuous mesh, contrasting with their introduction of surface cuts to accommodate human cerebellar anatomy, leading to a discontinuous flatmap.

Our flatmap enabled volume decomposition of the rat somatosensory cortex into non-overlapping hexagonal subvolumes of approximately equal size. These subvolumes can be used to group spatially distributed data, and perform statistical analyses that could reveal functional gradients along the horizontal axes. A similar approach was applied to the mouse pre-frontal cortex (PFC) in L. Gao et al., 2022 to analyze the modular structure of intra-PFC connectivity. Furthermore, in the context of data-driven modeling, subvolumes can be extracted to run simulations of smaller parts of a larger model Isbister et al., 2023 or used to characterize connectivity across scales Reimann, Bolaños-Puchet, et al., 2022.

Despite the numerous studies in rodent barrel cortex, previous efforts on barrel segmentation have been made only in rat (Egger et al., 2012), and existing mouse atlases lack annotations of individual barrels. We used our flatmap to derive annotations of barrels and their associated barrel columns with some notable properties. Firstly, they match previous accounts of barrels (Welker and Woolsey, 1974; Lefort et al., 2009) in terms of shapes and sizes. Secondly, they are defined in the standardized CCFv3 space, which enables the integration and analysis of datasets with single-barrel specificity. Finally, the barrel columns are non-overlapping and follow the curvature of the cortex, resulting in a more biologically accurate representation than would be given by intersecting straight cylinders.

Data visualization in flat space is similar to the staining of mechanically flattened tangential sections of cortex. This classic technique has been applied even in a per-layer fashion in previous work (Fig. 13 of Paperna and Malach, 1991, compare to Fig. 7C). However, unlike mechanical flattening, where “angular parts of the cortex undergo a certain distortion, which may lead to misinterpretation of the location of labeled cells” (Paperna and Malach, 1991), our approach avoids this curvature bias by taking into account the local orientation at each point. Another advantage of digital methods like ours over mechanical flattening is the ease of visualization of data from different individuals registered to the same reference space.

Topographical mapping in flat space (Fig. 7A) has also been previously reported. For example, a flatmap of axonal projections between entorhinal cortex and dentate gyrus of rat (Dolorfo and Amaral, 1998) was used to define connections in a computational model of hippocampus (Hendrickson et al., 2016). Flatmaps are also a key component of an algorithm used to predict a long-range micro-connectome of the whole mouse neocortex (Reimann, Gevaert, et al., 2019).

Notwithstanding the successes of our method, it has some of limitations. Notably, it is sensitive to edge artifacts. For example, in both atlases used in our results, some of the voxels at the edges of the source volume were not eligible. Since their numbers were very small compared to the total number of voxels, we did not try to fix them, but they still represent locations where our method failed to assign three-dimensional coordinates. These edge artifacts are usually a consequence of discontinuities in the relative depth or orientation fields, or of noise present in the atlas, and thus can in general be diminished by using a smoother atlas. Even when that is not a possibility, one strategy to mitigate edge artifacts could be to take a larger source volume, when possible, to effectively displace the edges away from the regions of interest. Alternatively, one could adopt a weaker eligibility criterion and keep all voxels whose streamlines reach the projection surface, even if these do not span the full depth. Ultimately, the missing coordinates could be filled in by extrapolation from nearby voxels.

Moreover, based on our flatmap metrics (Fig. 4F and Fig. 5E), locations with the largest non-uniformity and highest probability of non-continuity were also at the periphery of flat space. Beyond the above-mentioned impact of edge artifacts in the atlas, a possible cause for this may be the forced stretching of the projection mesh into a square (i.e., its boundary being made into straight lines regardless of its original shape) by the flattening algorithm. A mitigation strategy could be to extend the projection mesh to a square before flattening, so that the border would map more naturally to the unit square. There are many ways to do this, however, and care should be taken to not introduce additional artifacts.

One key parameter that can impact the quality of the flatmap is the depth *d*^∗^ at which the projection mesh is defined. Given how the flatmapping algorithm works, area-preservation and continuity of the flat space are best achieved at the selected depth, but as one moves away from *d*^∗^ these properties can degrade. This means the flatmap has highest fidelity of representation for voxels next to the projection mesh, and lower the farther away from it. In our results, we placed the projection mesh in the exact middle of the source volume (approximately at layer 4), but use cases that focus on more superficial or deeper layers could benefit from a different choice of *d*^∗^.

Another limitation stems from the use of streamlines to project voxels to the projection mesh. On the one hand, this ties the quality of the voxel projections to the quality of the orientation field, which may suffer from edge artifacts and discretization effects from limited resolution. On the other hand, while it is desirable that streamlines follow closely the geometry of a region, they are unavoidably subject to deformation depending on the relative size of the top and bottom shells at each location. For example, in slices of mouse prefrontal cortex the top of layer 1 can be seen to be more than four times wider than the bottom of layer 6 (Van De Werd et al., 2010); this geometrical narrowing underlies the “hotspots” visible in our flatmap metrics of mouse isocortex (Fig. 3.2E).

Beyond its limitations, our method offers flexibility in both implementation and applicability to diverse use cases. We have already shown that our flatmapping algorithm can be applied to data produced by other groups using other methods (Wang et al., 2020, see Results 3.2), as long as auxiliary atlases of local relative depth and orientation are available. If they are not, different approaches from ours can also be employed.

An alternative approach to generate relative depth and orientation fields is to solve the 3D heat equation inside the source volume, with “hot” top and “cold” bottom shells, and adiabatic sides (Jones, Buchbinder, and Aharon, 2000; Airey et al., 2005; Lerch et al., 2008); then, the solution itself can be used as the relative depth field and its gradient as the local orientation (see Results 3.2). Another similar, but simpler approach, consists in assigning the values 0 and 1 to the top and bottom shells, respectively, and then performing iterative 3D Gaussian blur to distribute these values smoothly throughout the source volume.

All of our, as well as these additional approaches, require the selection of top and bottom shells of the source volume. While this can be achieved by manual selection, this is a very time-consuming task that can benefit from (semi-)automated approaches. One possibility would be to use surface segmentation methods to partition the boundary mesh of the source volume into top, bottom and sides (Shamir, 2008; Nuvoli et al., 2021), and use this to guide voxel selection.

Regarding the flatmapping algorithm, while we have used specific algorithms in each step (and implementations thereof, see Supplementary material B), other algorithms can be used that achieve the same purpose. For example, there are many approaches in the field of computational geometry for level set extraction, surface reconstruction, mesh refinement and mesh flattening. The choice of alternatives by the user can serve to accommodate particular use cases or data. Also, while we have produced flatmaps that span large regions of cortex, for use cases that focus on single or small sets of cortical areas, it is possible to extract smaller parts of the projection mesh and use them to generate a flatmaps restricted to those areas.

Applications of flatmaps usually work at a certain pixel resolution, and our method allows the choice of the best one for each use case, using our metrics as guidelines. For visualization purposes, the primary concern is to avoid the presence of empty pre-images, i.e. holes in the flat view. Other use cases, such as volume decomposition and definition of barrel columns, work with shapes drawn on flat space and can be conducted at native resolution. Finally, use cases such as long-range axon visualization, require spatial continuity of the flat view, which may be attained at a lower resolution. This is especially crucial when gradients are considered in flat space (Guyonnet-Hencke and Reimann, 2023), as continuity guarantees they correspond to gradients in Cartesian space.

It is worth mentioning that it is also possible to use our method to generate flatmaps of brain regions having no laminar architecture, such as subcortical structures, as long as a principal axis can be meaningfully defined (see Supplementary material C for an example). A useful guideline to define the orientation of the principal axis is to consider the directions in which axons exit or enter the structure, as can be determined from 3D reconstructions of complete axons (e.g., Economo et al., 2016).

We think laminar coordinate systems offer new opportunities for data-driven research. For modeling, having a coordinate system adapted to the brain region being modeled enables capturing spatial features of biological complexity, such as the distribution of cell types or gradients of cell properties throughout the region. For example, the relative depth field can be used to define annotations of individual layers and to derive local distances to layer boundaries; together with the orientation field, this information can be used to place reconstructed neuronal morphologies accurately in a brain region, both in terms of location and orientation, leading to realistic anatomical models (e.g., Reimann, Bolaños-Puchet, et al., 2022).

Furthermore, our new flatmap-derived annotations of individual barrels and barrel columns provide a reference framework for future studies in barrel cortex, such as anatomical quantification and connectivity tracing (cortico-cortical and thalamocortical), as well as data-driven modeling.

In conclusion, we strongly believe that open and user-customizable methods to generate laminar coordinate systems and flatmaps are an important asset for the neuroscience community. Our contribution in this direction provides a basis for additional improvements allowing future neuroscience studies to benefit from these tools and approaches.

## Code and data availability

Enhanced atlases and flatmaps of rodent cortex are available in a public repository under the CC BY license: https://doi.org/10.5281/zenodo.8165004.

Code implementing the flatmapping algorithm and the definition of barrel cortex annotations is available in a public repository under a free software license: https://github.com/BlueBrain/atlas-enhancement.

## Acknowledgements

This study was supported by funding to the Blue Brain Project, a research center of the École polytechnique fédérale de Lausanne (EPFL), from the Swiss government’s ETH Board of the Swiss Federal Institutes of Technology.

We’d like to thank Karin Holm for useful comments and copy editing.

## Supplementary material

### A. Atlas of rat somatosensory cortex

This atlas was built in-house as the first step of an atlas-based model building effort that culminated in a large-scale biophysically detailed model of somatosensory cortex of juvenile rat (Reimann, Bolaños-Puchet, et al., 2022).

The starting point for the atlas was the digital component of Paxinos & Watson’s printed atlas of the adult rat brain (CD-ROM distributed with Paxinos and Watson, 2007), consisting of vector drawings of coronal slices with annotated region outlines. These individual slices were aligned, rasterized and interpolated, then combined to create an isotropic voxelized atlas (at 40 *µ*m resolution) with region labels corresponding to the region and acronym index in the book. The somatosensory regions in one hemisphere were then extracted and hierarchically smoothed to create an atlas of rat somatosensory cortex. This atlas was down-scaled by a factor of 2082*/*2150 = 0.96837, based on cortical thickness of the S1HL region, to approximate the size of a juvenile (P14) rat brain.

The final atlas consists of an array of 409 × 608 × 286 voxels with size 38.7348 *µ*m with 10 annotated regions, including primary somatosensory representations of barrel field (S1BF), front limb (S1FL), hind limb (S1HL), jaw (S1J), shoulder (S1Sh), trunk (S1Tr), upper lip (S1ULp), dysgranular zone (S1DZ) and dysgranular oral zone (S1DZO); as well as secondary somatosensory region (S2) and an unspecified primary somatosensory region (S1).

### B. Implementation details

#### General

- All code was run under GNU/Linux on the BB5 supercomputer hosted at the Swiss National Supercomputing Centre (CSCS).
- The workflow orchestrating all steps of the algorithm is written as a GNU Makefile.
- Handling of atlas volumes in NRRD format is performed in Python using `voxcell` (Povolotsky et al., 2023).

#### Flatmapping stage I: Flat mesh generation

- Extraction of the point cloud of voxels approximately at relative depth *d*^∗^ is implemented in Python using the Euclidean Distance Transform (EDT) algorithm in SciPy.
- Reconstruction of the projection mesh is implemented in C++ using the Scale-Space Surface Reconstruction package from the Computational Geometry Algorithms Library (CGAL, Lankveld, 2023).
- Refinement of the projection mesh by uniform subdivision is performed with GMSH (Geuzaine and Remacle, 2009).
- Flattening of the projection mesh is performed using the (Iterative) Authalic Mapping algorithm from the Triangulated Surface Mesh Parameterization CGAL package (Saboret et al., 2023).

#### Flatmapping stage II: Voxel projection

- Integration of streamlines is implemented in C using the numerical ODE integrator with explicit embedded Runge-Kutta (2,3) method from the GNU Scientific Library (GSL, Galassi, 2009).
- Parametrization of streamlines is implemented in C using linear spline interpolation from GSL.
- Projection of voxel centers to the projection surface is implemented in C using the Brent-Dekker one-dimensional root finding algorithm from GSL.
- Distributed parallel computation of voxel projections is performed with the help of GNU Parallel (Tange, 2011).

#### Flatmapping stage III: Flatmap generation

- Mapping of streamline intersections to mesh points is implemented in C++ using Nearest Neighbor search from the dD Spatial Search CGAL package (Tangelder and Fabri, 2023).
- Discretization of the flatmap is performed in Python using NumPy.

#### Applications

- Plotting of flat views is performed in Python using Datashader.

### C. Flatmapping of non-layered brain structures

Our flatmapping method can also be applied to non-layered brain regions, as long as a principal axis can be defined that is meaningful for the geometry of the structure under study.

Here we provide as an example the generation of a flatmap for the dorsal striatum of mouse. This structure has no cytoarchitectonically defined layers, but it has a smooth organic shape with a meniscus cross-section (similar to a cupped hand) that can be approximated by an ellipsoid. The principal axis is then defined as the normal vector to the surface of the ellipsoid, and the flatmap axes go along the surface of the ellipsoid.

In detail:

1. We extract the voxels belonging to the dorsal striatum (STRd) region from the Allen Mouse Brain Atlas CCFv3 at 25 um resolution.
2. We fit a plane to the point cloud defined by the voxel centers, in order to get an idea of the “facing” direction of the structure.
3. We fit a sphere to the point cloud on the concave side, with center along the line defined by the plane normal.
4. Using the position and radius of the sphere as initial condition, we fit an ellipsoid using the sum of orthogonal distances as the cost function with the COBYLA optimization algorithm (NLopt library). The resulting ellipsoid passes through the “middle” of the point cloud, similar to the choice of isosurface with relative depth 0.5 in the main text.
5. We compute the bounding box of the point cloud and intersect this with the ellipsoid to obtain a patch of ellipsoidal surface that we use as projection mesh.
6. We compute the projections of voxel centers as orthogonal projection to the ellipsoid, that is, streamlines are assumed to be straight line segments having the orientation of the surface normals.
7. We compute relative depth by considering the normalized distance to the ellipsoid surface, taking into account which side of the surface the point is in, so we obtain a continuous value in [0,1].
8. The rest of the algorithm proceeds as usual, with authalic flattening of the projection mesh and finding nearest neighbors between intersection points and mesh vertices.

The resulting flatmap provides coordinates along the “face” of the dorsal striatum, and allows a decomposition of its volume into uniform subvolumes. A similar approach can be applied to any brain structure with a similar shape, e.g. reticular nucleus of thalamus, etc.

**Supplementary Figure 1:**
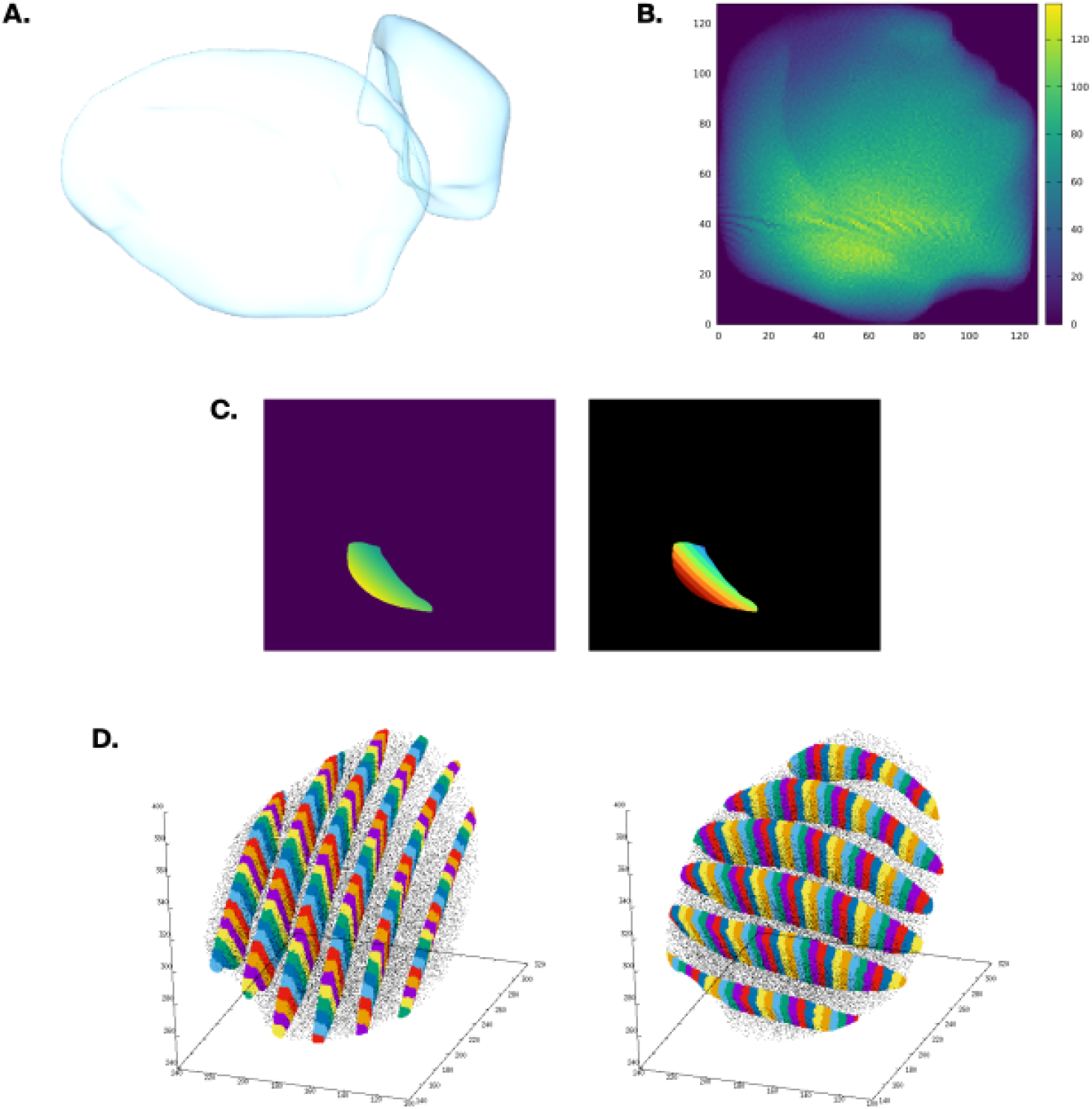
Flatmapping of non-layered brain structures: dorsal striatum. **A**. 3D meshes of bilateral dorsal striatum, showing “face” of structure and lateral aspect. **B**. Pre-image size heatmap. Note how the edges of the flatmap do not coincide with the edges of the square due to how the projection mesh was chosen. **C**. Slice through the relative depth field, showing continuous variation (left) and discrete levels to exhibit ellipsoidal shape of isosurfaces (right). **D**. Decomposition of the volume of dorsal striatium into square columns. Note how the columns curve following the normals of the ellipsoidal surface and span the full depth.

### D. Radial volume distribution of a truncated square pyramid

When computing the width of the pre-image of a pixel, we perform a fit based on the radial volume distribution of the pre-image, using the best-fit line through the centroid as axis. We fit the two radii of a truncated square pyramid based on the analytical expression for the normalized radial volume distribution, which we describe here.

Essentially, we are computing the volume of the intersection between a truncated square pyramid with radii *r*_0_, *r*_1_ (*r*_0_ ≤ *r*_1_) and height *H*, and a cylinder of radius *r* with the same axis. To compute this volume as a function of *r*, we take the integral along height of the cross-sectional area of the intersection.

The cross-section at any point corresponds to the intersection between a circle of radius *r* and a square of radius *a* concentric to it, which has an area (Fig. S2 A):

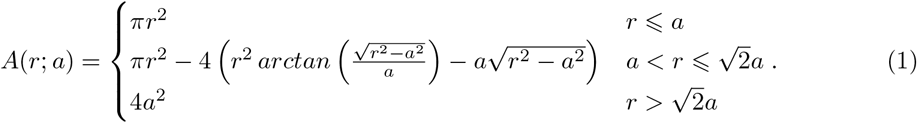

In the case of the truncated square pyramid, *a* depends linearly on height:

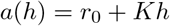

with 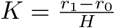, so we can write the volume integral as:

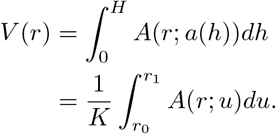

Now, since *A*(*r*; *u*) is defined piecewise, this integral ends up split into intervals based on the range of *r* relative to *u*. In any case, the result is a combination of the definite integrals of the three cases in (1), which are given by (Fig. S2 B):

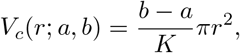

the volume of a cylinder of radius *r* and height 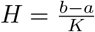,

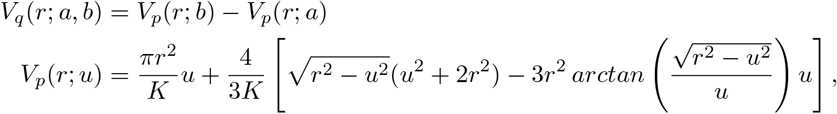

the volume of a proper intersection between a cylinder of radius *r* and a truncated square pyramid with radii *a, b* (*b* ≥ *a*), and

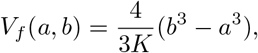

the full volume of a truncated square pyramid with radii *a, b* (*b* ≥ *a*) and height 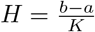.

The following special cases of *V*_*p*_(*r*; *u*) are useful for numerical evaluation:

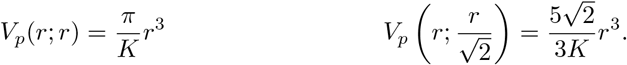

In terms of these integrals, and considering the different ranges of *r*, we write the radial volume distribution of a truncated square pyramid with radii *r*_0_, *r*_1_ (*r*_0_ ≤ *r*_1_) and height 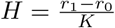 as:

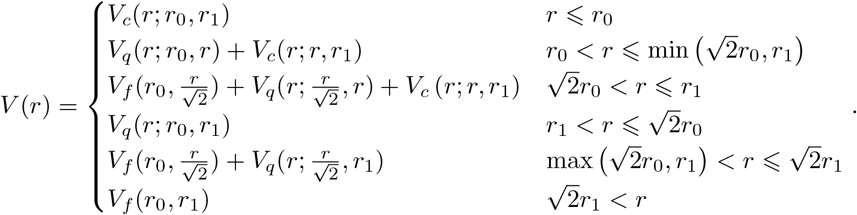

The third and fourth cases are exclusive to one another, with the third one occurring only when 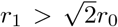 and the fourth one in the opposite case. When *r*_1_ → *r*_0_, the function tends to the volume of the square prism.

By considering the quotient with the total volume, i.e. *V* (*r*)*/V*_*f*_ (*r*_0_, *r*_1_), we obtain the volume fraction, a function with values in [0, 1] that only depends on *r*_0_ and *r*_1_, but not on *H* (since all terms have a factor of 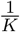 that cancels out when taking the quotient). This function can be readily implemented numerically and fitted using a least squares approach to the empirical cumulative distribution function (CDF) of the data (Fig. S2 C).

**Supplementary Figure 2:**
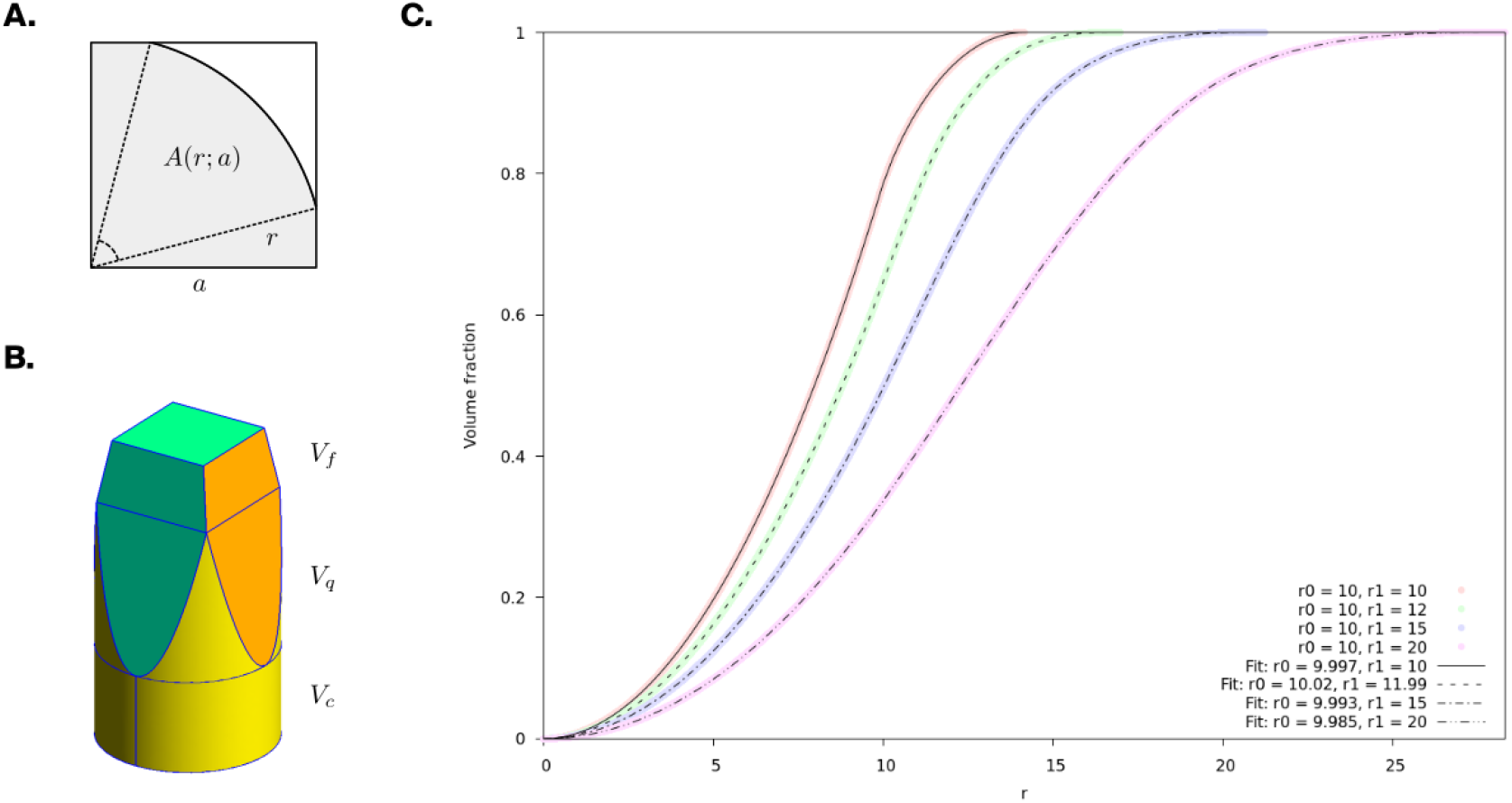
Radial volume distribution of a truncated square pyramid. **A**. Geometry of the intersection between a circle of radius *r* and a square of radius *a*, from which *A*(*r*; *a*) in Equation (1) can be derived using basic trigonometry. **B**. 3D schematic of the volumes described by the three cases of the volume integral: truncated square pyramid (*V*_*f*_, top), cylinder (*V*_*c*_, bottom), and their intersection (*V*_*q*_, middle). **C**. Least squares fits of the normalized radial volume distribution to the empirical CDF of data generated by random sampling of a perfect truncated square pyramid with given radii. An excellent fit is obtained in all cases: 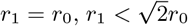 and 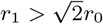, providing empirical evidence of the applicability of our analytical derivation.

## Notes

https://zenodo.org/doi/10.5281/zenodo.8165004

### Competing Interest Statement

The authors have declared no competing interest.

### Summary of Updates

Title updated. Author list updated. Revised writing and copy editing in all sections. Improved most figures. Introduced new metrics in Results 3.3.

